## Supplementary Information for "Engineering an artificial catch bond using mechanical anisotropy"

#### Supplementary figures

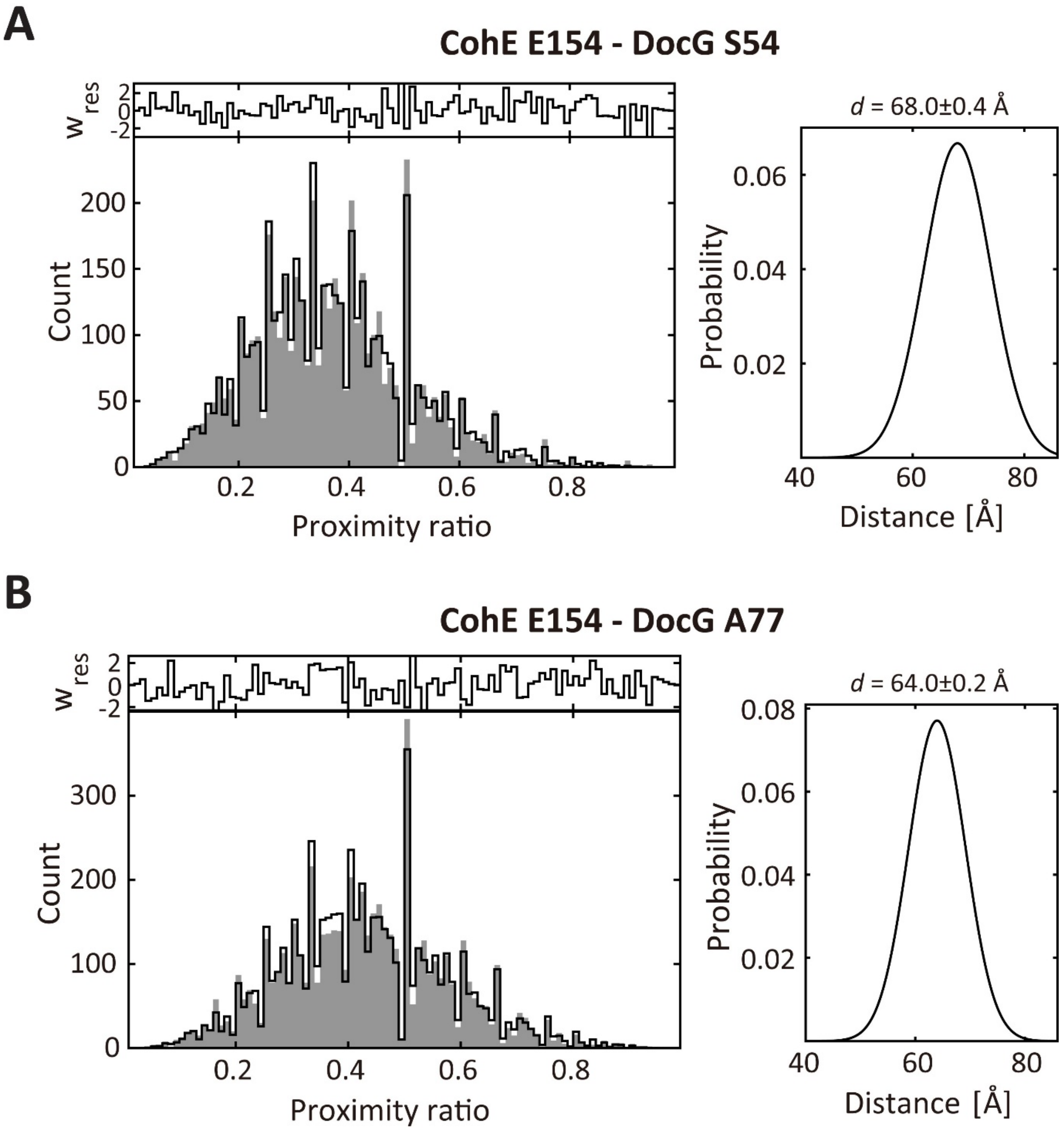


**Figure S1. Photon density analysis (PDA) results.** Left panel: simulated proximity ratio histograms of CohE E154 - DocG S54 measurement **(A)** and CohE E154 - DocG A77 measurement **(B)**. Right panel: probability density distribution of simulated donor-acceptor distance between CohE E154 and DocG S54 **(A)** and CohE E154 and DocG A77 **(B)**.

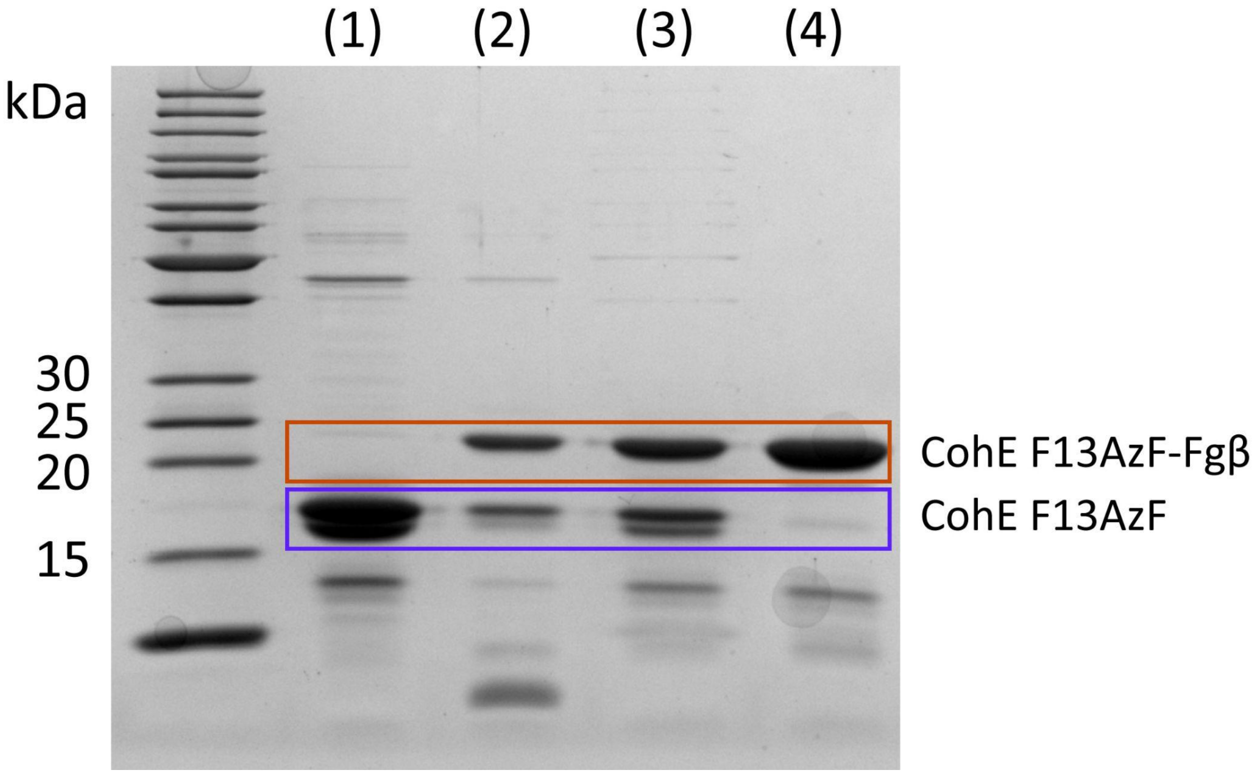


**Figure S2. SDS-PAGE showing conjugation of Fgβ to Coh F13AzF mutant.** The Coh F13AzF mutant (Lane 1) was conjugated with Fgβ peptide. The reaction product (Lane 2) was subsequently purified using a size-exclusion column to remove the excess Fgβ peptide (Lane 3), and a Strep-trap column to remove unconjugated Coh F13AzF (Lane 4).

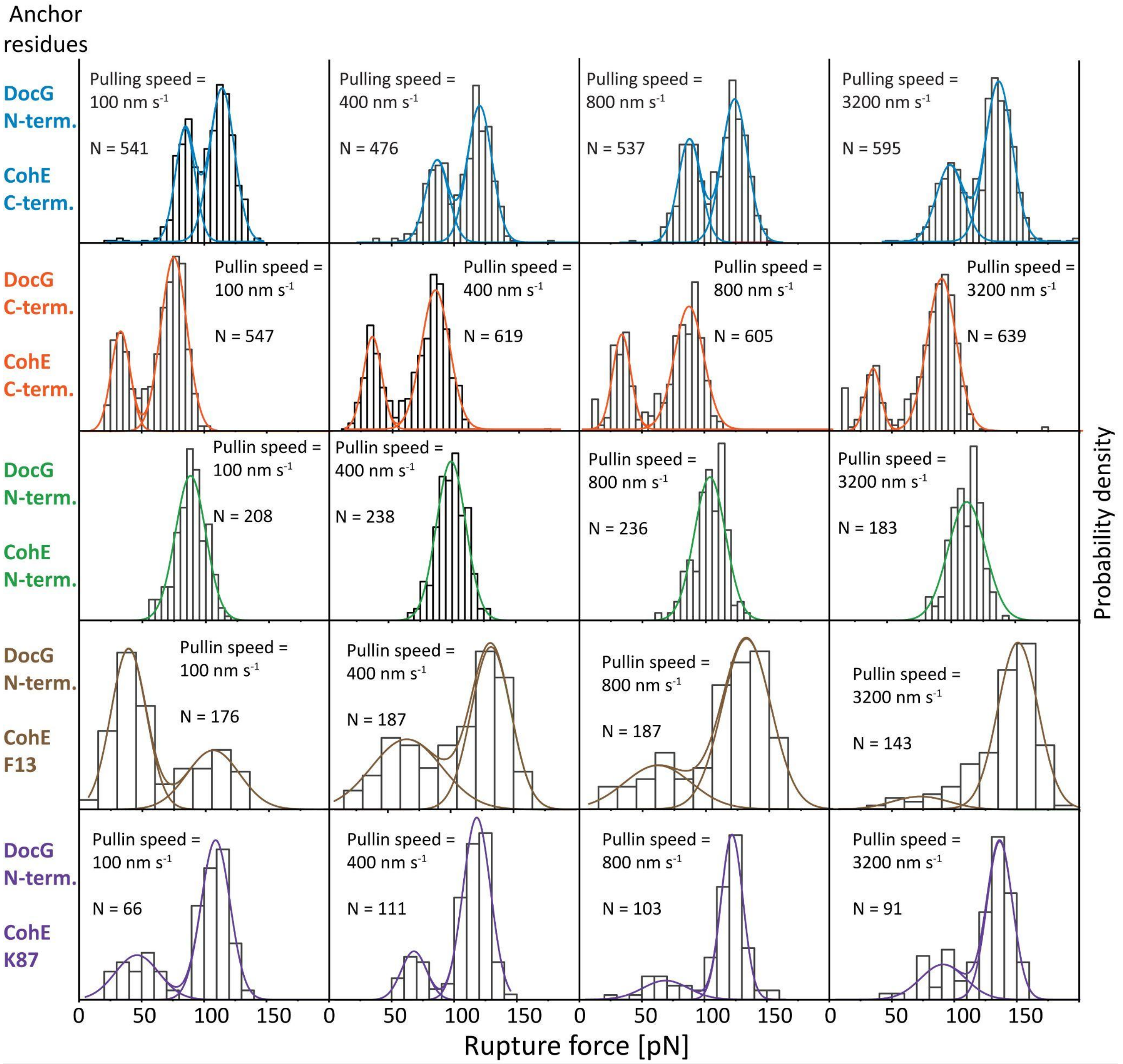


**Figure S3 Rupture force histograms measured at different anchor geometries and different pulling speeds.** The histograms were fitted with one (CohE N-terminus and DocG N-terminus anchor geometry) or two (other anchor geometries) Gaussian peaks.

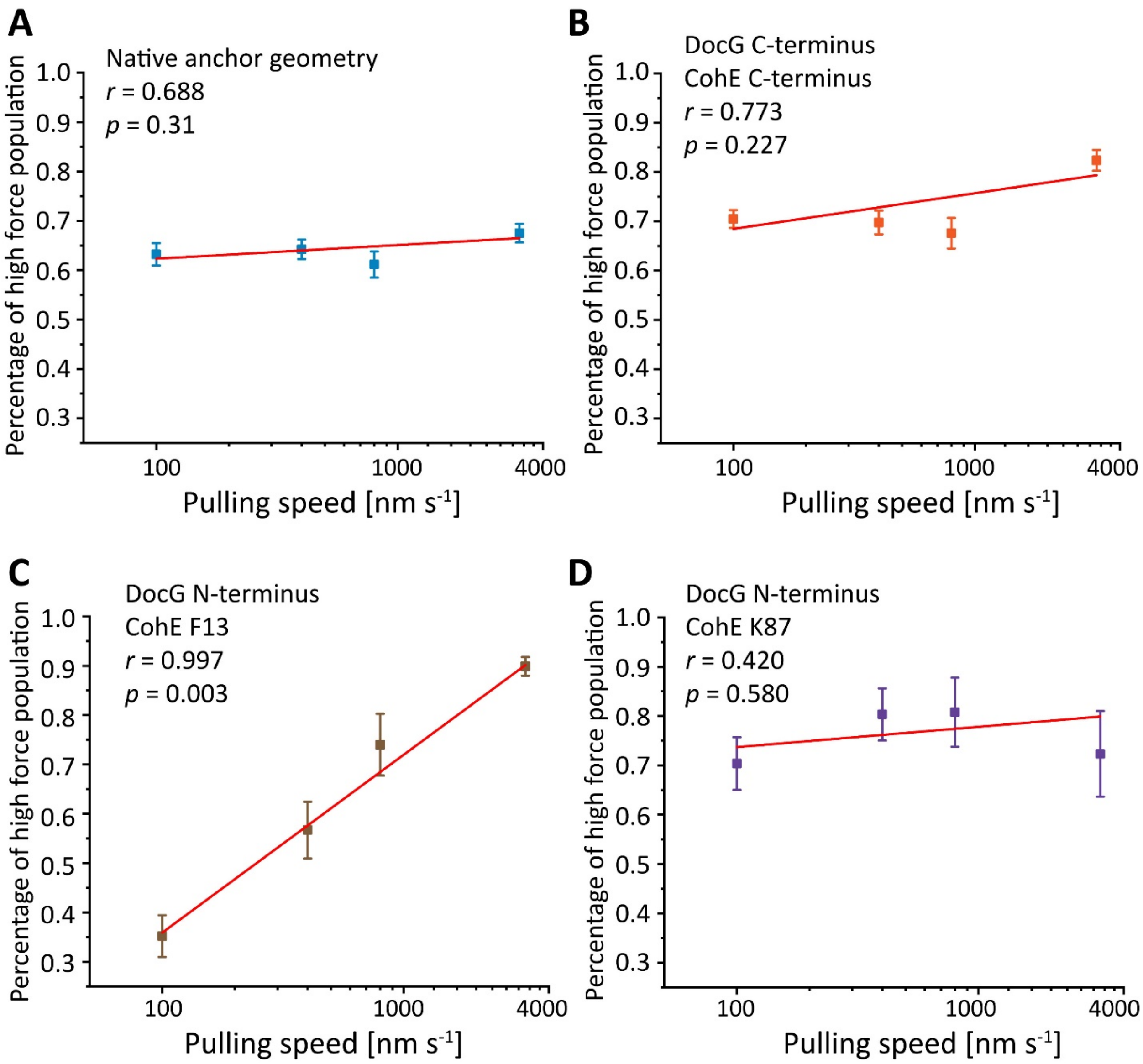


**Figure S4 Linear regression between the percentage of high force population and the logarithm of pulling speed.** For each anchor geometry, the prevalence of the high force population measured at different pulling speeds are plotted against the logarithm of the pulling speed and fitted linearly. The Pearson's correlation coefficients and the *p*-values are shown for each anchor geometry and summarized in Table S2.

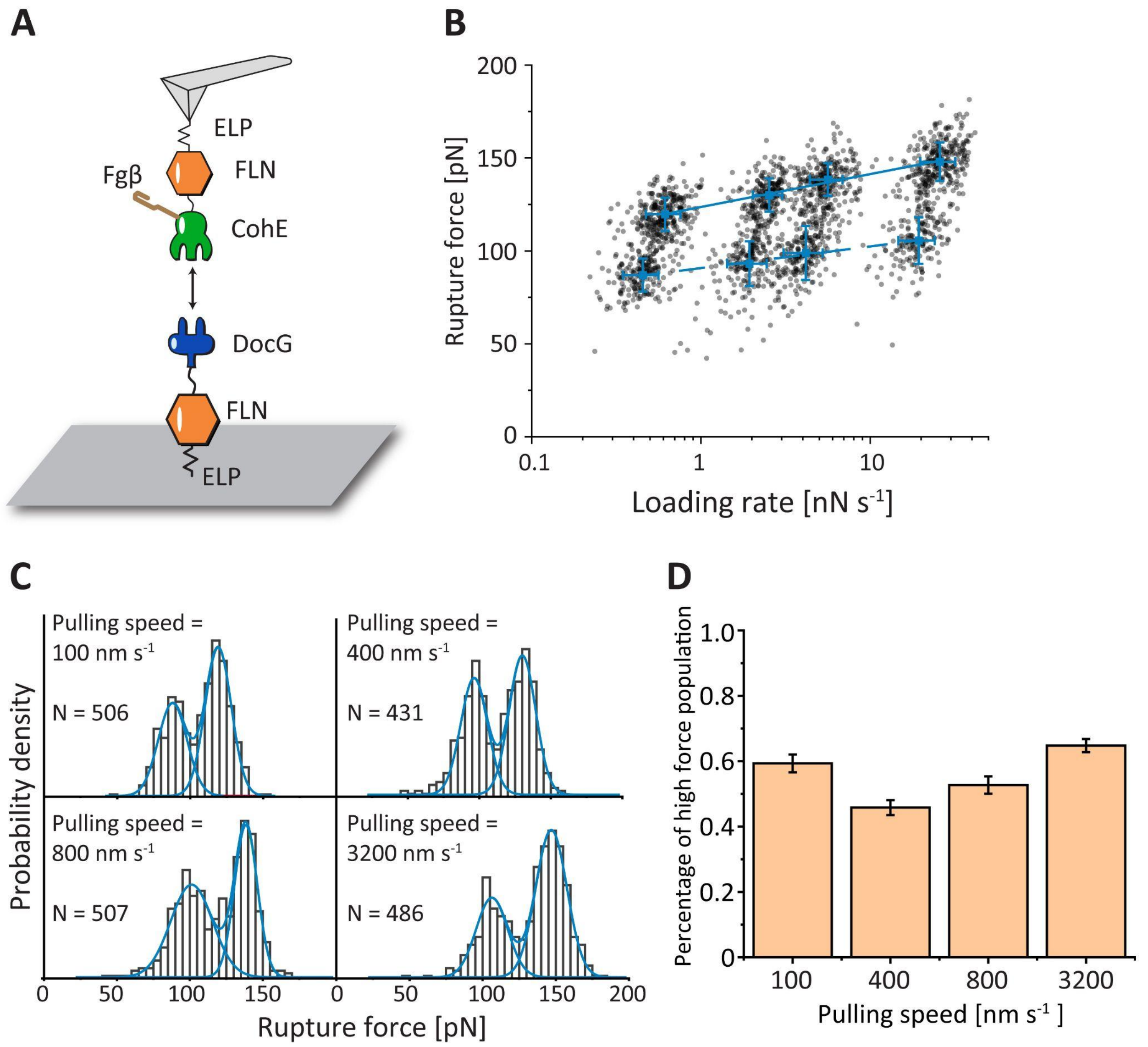


**Figure S5 Native anchor geometry measurement of DocG:CohE (F13-Fgβ). A:** AFM-SMFS experimental setup. The residue F13 of CohE in CohE-FLN-ELP-ybbr construct was replaced by azido-phenylalanine and conjugated with Fgβ peptide, and the construct was immobilized on the AFM tip. ybbr-ELP-FLN-DocG construct was immobilized on the glass surface. The rupture force of DocE:CohG (F13-Fgβ) complex was measured at the native anchor geometry and at different pulling speeds. **B:** The force-loading rate plot of DocG:CohE (F13-Fgβ). The average rupture forces of the high force and low force pathways measured at four different pulling speeds were linearly fitted against loading rate to extract energy landscape parameters. **C:** Rupture force histograms of DocG:CohE (F13-Fgβ) at different pulling speeds. **D:** Fraction of high force pathway at different pulling speeds.

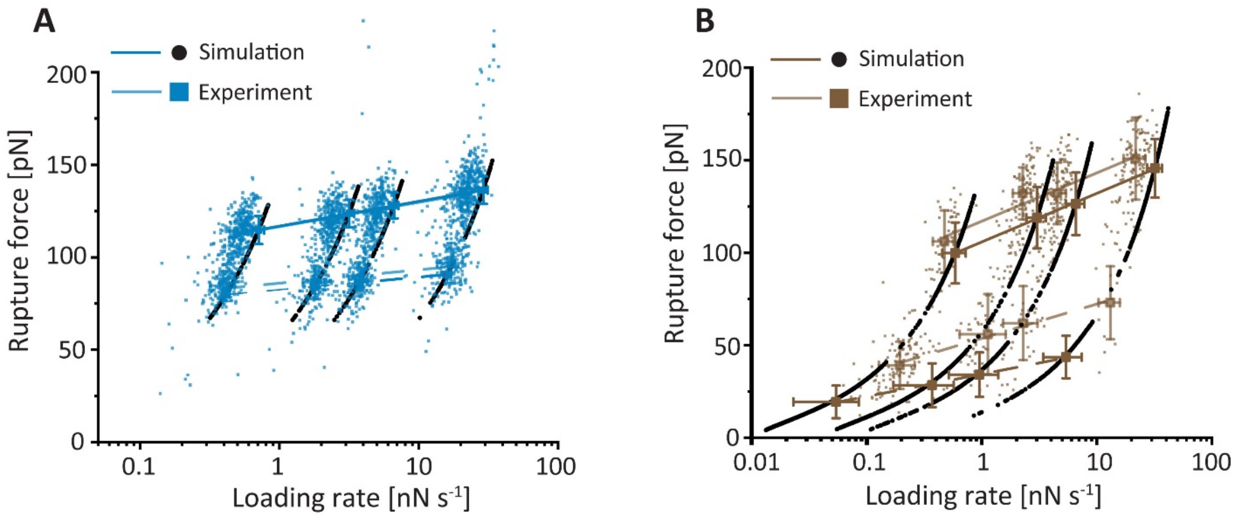


**Figure S6 Overlay of force-loading rate plots of experimental and simulation results.** The Monte Carlo simulation results for the force-loading rate relationship at the native (**A**) and catch (**B**) anchor geometries are overlayed with the corresponding experimental results. The simulation results are shown in black dots and the experimental data are shown in blue (native anchor geometry) and brown (catch anchor geometry) squares. The Bell-Evans fitting results are shown in solid line (high-force population) and dashed line (low-force population).

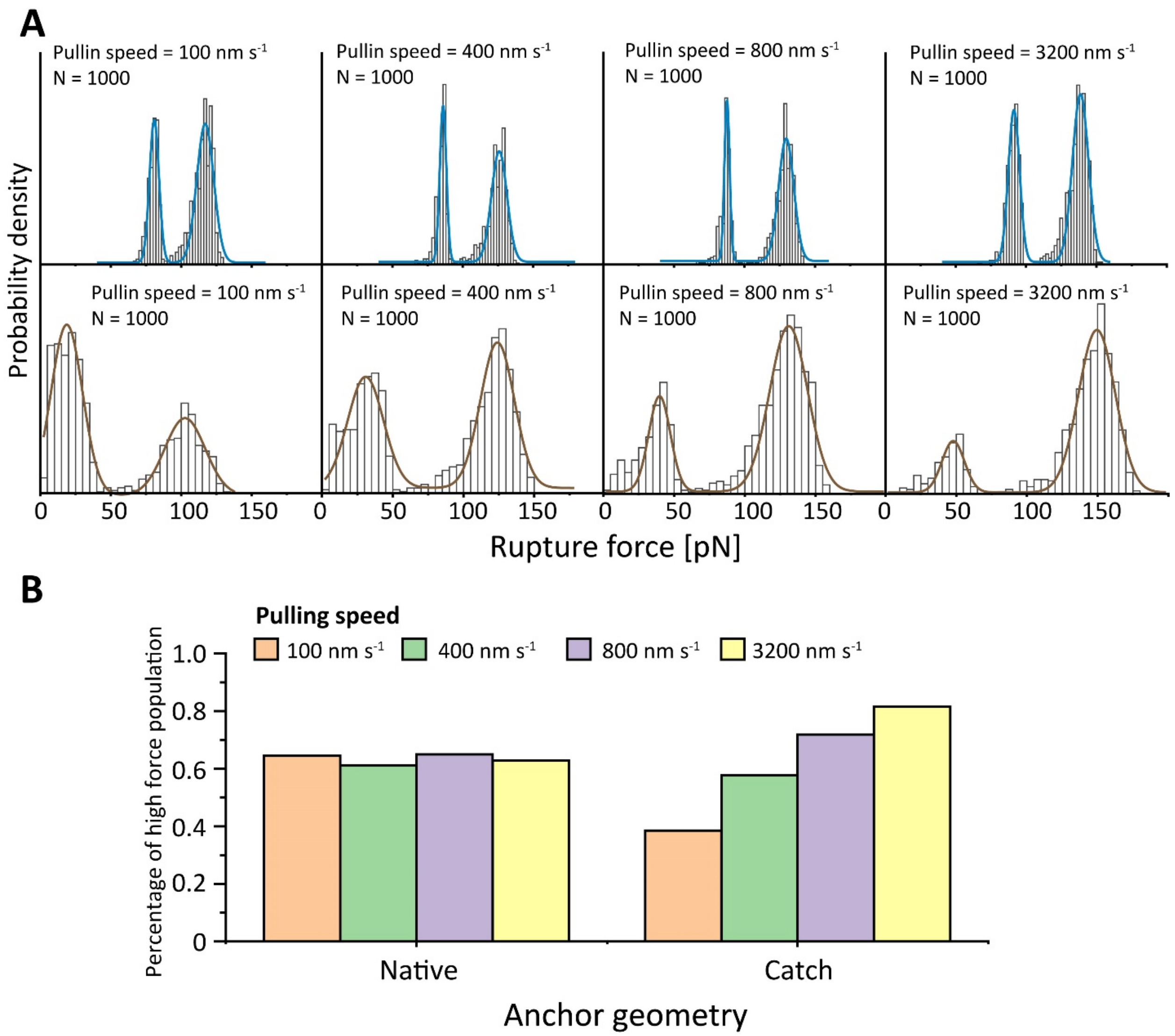


**Figure S7 Monte Carlo simulation rupture force histograms. A:** Rupture force histograms of DocG:CohE at native anchor geometry (upper panels) and catch anchor geometry (anchored at residue F13 of CohE, lower panels). Each histogram was fitted in two Gaussian peaks. **B:** The prevalence of high rupture force population at different pulling speeds in Monte Carlo simulation. At the catch anchor geometry, the prevalence of high force population increases with increasing pulling speed ($p < 0.05$, see **Table S2**), which is in contrast to the native anchor geometry.

#### Supplementary tables

**Table S1 Equilibrium affinity of the Doc:Coh complex**

| | Wild-type CohE | CohE F13AzF-Fgβ | CohE K87AzF-Fgβ |
|---|---|---|---|
| Dissociation constant [nM] | 14±4 | 40±8 | 19±10 |

**Table S2 Pearson's correlation coefficients between logarithm of pulling speed and prevalence of high force unbinding pathway**

| **Anchor geometry** | **Pearson's correlation coefficient *r*** | ***p*-value** |
|---|---|---|
| Native (experiment) | 0.688 | 0.31 |
| Native (Monte Carlo simulation) | -0.196 | 0.804 |
| DocG C-terminus CohE C-terminus (experiment) | 0.773 | 0.227 |
| DocG N-terminus CohE N-terminus (experiment) | N/A | N/A |
| DocG N-terminus CohE F13 (experiment) | 0.997 | 0.003 |
| DocG N-terminus CohE F13 (Monte Carlo simulation) | 0.982 | 0.02 |
| DocG N-terminus CohE K87 (experiment) | 0.420 | 0.580 |

**Table S3 Unbinding energy landscape parameters calculated from Monte Carlo simulations**

| Anchor geometry | Pathway | log($k_0$) | $\Delta x^{\ddagger}$ [nm] |
|---|---|---|---|
| Native (slip) | High force | -6.5±0.1 | 0.720±0.009 |
| | Low force | -9.9±0.2 | 1.42±0.02 |
| DocG N-terminus CohE F13 (catch) | High force | -1.80±0.09 | 0.368±0.008 |
| | Low force | -0.6±0.1 | 0.80±0.03 |

### Supplementary notes

#### Supplementary note 1 Amino acid sequences

Color code:

ybbr tag, His tag, ELP linker, FLN, DocG, CohE, AzF

**ybbr-His-ELP-FLN-DocG**

MGTDSLEFIASKLAHHHHHHWGSGHGVGVPGMGVPGVGVPGVGVPGVGVPGVGVPGVGVPG
VGVPGVGVPGEGVPGEGVPGVGVPGMGVPGVGVPGVGVPGVGVPGVGVPGVGVPGVGVPGV
GVPGEGVPGEGVPGVGVPGMGVPGVGVPGVGVPGVGVPGVGVPGVGVPGVGVPGVGVPGEG
VPGEGVPGWPSGSADPEKSYAEGPGLDGGESFQPSKFKIHAVDPDGVHRTDGGDGFVVTIEGPA
PVDPVMVDNGDGTYDVEFEPKEAGDYVINLTLDGDNVNGFPKTVTVKPAPGSGSGSGSGVGDS
LLRGDVDLDGDVDVADAVAVLQASAEQMVTGENPLSKDARFGADVNDDSRVDVSDAVLILQY
SSMKIANPDADWDDLLG

**CohE-FLN-ELP-His-ybbr**

ADGAAKLSMDQKFAEPGETVEIALNLENFDASWTGLEFLVNYDPKLEVALDGAGDIDYSYGDAI
GAMGKKISVGGAISKDLTADGLKGFAFAWGTATAISGNGQLGVFKFTVPADAQPGDEFPVNLTV
NVGSFIDANKENIPFETVNGWIKIKEEGSGSGSGSADPEKSYAEGPGLDGGECFQPSKFKIHAVDP
DGVHRTDGGDGFVVTIEGPAPVDPVMVDNGDGTYDVEFEPKEAGDYVINLTLDGDNVNGFPKT
VTVKPAPGSGSGSHGVGVPGMGVPGVGVPGVGVPGVGVPGVGVPGVGVPGVGVPGVGVPGEG
VPGEGVPGVGVPGMGVPGVGVPGVGVPGVGVPGVGVPGVGVPGVGVPGVGVPGEGVPGEGVP
GVGVPGMGVPGVGVPGVGVPGVGVPGVGVPGVGVPGVGVPGVGVPGEGVPGEGVPGWRGHH
HHHHGSDSLEFIASKLA

**DocG-FLN-ELP-His-ybbr**

GVGDSLLRGDVDLDGDVDVADAVAVLQASAEQMVTGENPLSKDARFGADVNDDSRVDVSDA
VLILQYSSMKIANPDADWDDLLGGSGSGSADPEKSYAEGPGLDGGECFQPSKFKIHAVDPDGVH
RTDGGDGFVVTIEGPAPVDPVMVDNGDGTYDVEFEPKEAGDYVINLTLDGDNVNGFPKTVTVK
PAPGSGSGSHGVGVPGMGVPGVGVPGVGVPGVGVPGVGVPGVGVPGVGVPGVGVPGEGVPGE
GVPGVGVPGMGVPGVGVPGVGVPGVGVPGVGVPGVGVPGVGVPGVGVPGEGVPGEGVPGVG
VPGMGVPGVGVPGVGVPGVGVPGVGVPGVGVPGVGVPGVGVPGEGVPGEGVPGWRGHHHHH
HGSDSLEFIASKLA

**ybbr-His-ELP-FLN-CohE**

MGTDSLEFIASKLAHHHHHHWGSGHGVGVPGMGVPGVGVPGVGVPGVGVPGVGVPGVGVPG
VGVPGVGVPGEGVPGEGVPGVGVPGMGVPGVGVPGVGVPGVGVPGVGVPGVGVPGVGVPGV
GVPGEGVPGEGVPGVGVPGMGVPGVGVPGVGVPGVGVPGVGVPGVGVPGVGVPGVGVPGEG
VPGEGVPGWPSGSADPEKSYAEGPGLDGGESFQPSKFKIHAVDPDGVHRTDGGDGFVVTIEGPA
PVDPVMVDNGDGTYDVEFEPKEAGDYVINLTLDGDNVNGFPKTVTVKPAPGSGSADGAAKLSM
DQKFAEPGETVEIALNLENFDASWTGLEFLVNYDPKLEVALDGAGDIDYSYGDAIGAMGKKISV
GGAISKDLTADGLKGFAFAWGTATAISGNGQLGVFKFTVPADAQPGDEFPVNLTVNVGSFIDAN
KENIPFETVNGWIKIKEE

**CohE (F13AzF)-FLN-ELP-His-ybbr**

ADGAAKLSMDQKAzFAEPGETVEIALNLENFDASWTGLEFLVNYDPKLEVALDGAGDIDYSYG
DAIGAMGKKISVGGAISKDLTADGLKGFAFAWGTATAISGNGQLGVFKFTVPADAQPGDEFPVN
LTVNVGSFIDANKENIPFETVNGWIKIKEEGSGSGSGSADPEKSYAEGPGLDGGECFQPSKFKIHA
VDPDGVHRTDGGDGFVVTIEGPAPVDPVMVDNGDGTYDVEFEPKEAGDYVINLTLDGDNVNGF
PKTVTVKPAPGSGSGSHGVGVPGMGVPGVGVPGVGVPGVGVPGVGVPGVGVPGVGVPGVGVP
GEGVPGEGVPGVGVPGMGVPGVGVPGVGVPGVGVPGVGVPGVGVPGVGVPGVGVPGEGVPGE
GVPGVGVPGMGVPGVGVPGVGVPGVGVPGVGVPGVGVPGVGVPGVGVPGEGVPGEGVPGWR
GHHHHHHGSDSLEFIASKLA

**CohE (F13AzF)-FLN-ELP-His-ybbr**

ADGAAKLSMDQKAzFAEPGETVEIALNLENFDASWTGLEFLVNYDPKLEVALDGAGDIDYSYG
DAIGAMGKKISVGGAISKDLTADGLKGFAFAWGTATAISGNGQLGVFKFTVPADAQPGDEFPVN
LTVNVGSFIDANKENIPFETVNGWIKIKEEGSGSGSHHHHHH

**CohE (K87AzF)-FLN-ELP-His-ybbr**

ADGAAKLSMDQKFAEPGETVEIALNLENFDASWTGLEFLVNYDPKLEVALDGAGDIDYSYGDAI
GAMGKKISVGGAISKDLTADGLAzFGFAFAWGTATAISGNGQLGVFKFTVPADAQPGDEFPVNL
TVNVGSFIDANKENIPFETVNGWIKIKEEGSGSGSHHHHHH
